# Post-transcriptional cross- and auto-regulation buffer expression of the human RNA helicases *DDX3X* and *DDX3Y*

**DOI:** 10.1101/2024.07.08.602613

**Authors:** Shruthi Rengarajan, Jason Derks, Daniel W. Bellott, Nikolai Slavov, David C. Page

## Abstract

The Y-linked gene *DDX3Y* and its X-linked homolog *DDX3X* survived the evolution of the human sex chromosomes from ordinary autosomes. *DDX3X* encodes a multi-functional RNA helicase, with mutations causing developmental disorders and cancers. We find that, among X-linked genes with surviving Y homologs, *DDX3X* is extraordinarily dosage-sensitive. Studying cells of individuals with sex chromosome aneuploidy, we observe that when the number of Y chromosomes increases, *DDX3X* transcript levels fall; conversely, when the number of X chromosomes increases, *DDX3Y* transcript levels fall. In 46,XY cells, CRISPRi knockdown of either *DDX3X* or *DDX3Y* causes transcript levels of the homologous gene to rise. In 46,XX cells, chemical inhibition of DDX3X protein activity elicits an increase in *DDX3X* transcript levels. Thus, perturbation of either *DDX3X* or *DDX3Y* expression is buffered – by negative cross-regulation of *DDX3X* and *DDX3Y* in 46,XY cells, and by negative auto-regulation of *DDX3X* in 46,XX cells. *DDX3X*-*DDX3Y* cross-regulation is mediated through mRNA destabilization – as shown by metabolic labeling of newly transcribed RNA – and buffers total levels of DDX3X and DDX3Y protein in human cells. We infer that post-transcriptional auto-regulation of the ancestral (autosomal) *DDX3* gene transmuted into auto- and cross-regulation of *DDX3X* and *DDX3Y* as these sex-linked genes evolved from ordinary alleles of their autosomal precursor.

## Introduction

*DDX3X* and *DDX3Y* are homologous but non-identical genes on the human X and Y chromosomes (Lahn and Page 1997). They encode pleiotropic RNA helicases implicated in multiple aspects of RNA metabolism, including splicing, export, stability, translation, and stress response (Soto-Rifo and Ohlmann 2013). *DDX3X* is widely conserved across eukaryotes, with orthologs in mammals, flies, worms, and yeast (Elbaum-Garfinkle et al. 2015; Johnstone et al. 2005; Sharma et al. 2017). Human *DDX3X* mutations are associated with several neurodevelopmental disorders and cancers (Snijders Blok et al. 2015; Valentin-Vega et al. 2016). *DDX3X* is expressed throughout the body from the “inactive” X chromosome (Xi) in females as well as from the “active” X chromosome (Xa) in males and females (Lahn and Page 1997; Tukiainen et al. 2017). Like *DDX3X*, its Y-chromosomal homolog *DDX3Y* is expressed in a wide array of human tissues (Godfrey et al. 2020), but studies of its organismal function have focused on roles in spermatogenesis (Ramathal et al. 2015). The X- and Y-encoded proteins are 91% identical at the amino acid level (Lahn and Page 1997). While they have significantly diverged in their N- and C-terminal regions, the RNA binding and helicase domains are largely conserved (Rosner and Rinkevich 2007). Early experiments showed that DDX3Y protein was functionally interchangeable with DDX3X *in vitro* (Sekiguchi et al. 2004). More recent work has shown that the proteins have partially overlapping functions, with similar effects on protein synthesis (Venkataramanan et al. 2021) but differing capacities for stress granule formation and translational repression (Shen et al. 2022; Venkataramanan et al. 2021).

*DDX3X* and *DDX3Y* constitute one of only 17 human X-Y gene pairs that survived the sex chromosomes’ evolution from ordinary autosomes (Lahn and Page, 1999; Skaletsky et al. 2003). While the human X chromosome retains 98% of the genes that were present on the ancestral autosomes, the Y chromosome retains only 3% of these genes (Bellott et al. 2014). Most of these surviving Y chromosome genes were preserved by natural selection to maintain the ancestral dosage of regulators of key cellular processes. Among this select group of X-linked genes with surviving Y homologs, we recently noticed a distinguishing feature of *DDX3X*: while the gene is robustly expressed from both Xa and Xi in human cells (and in this sense resembles other X-linked genes with surviving Y homologs), steady-state levels of *DDX3X* transcripts were only modestly higher in 46,XX cells than in 46,XY cells (San Roman et al. 2023), suggesting that *DDX3X* (and possibly *DDX3Y*) might be subject to dosage constraints and regulatory mechanisms not seen with other X-Y gene pairs. Accordingly, we decided to examine closely the dosage sensitivity and regulation of *DDX3X* and *DDX3Y*.

Here, we report that *DDX3X* and *DDX3Y* are extraordinarily dosage sensitive, even when compared with other human X-Y gene pairs. Their dosage is buffered by negative post-transcriptional cross-regulation of *DDX3X* and *DDX3Y* in 46,XY cells, while *DDX3X* is post-transcriptionally auto-regulated in 46,XX cells. We suggest that posttranscriptional cross-regulation of *DDX3X* and *DDX3Y* reflects the conservation in both genes of auto-regulatory mechanisms that governed their unitary autosomal precursor.

## Results

### *DDX3X* and *DDX3Y* are especially dosage-sensitive compared to genes with a similar evolutionary trajectory

We first asked if *DDX3X* and *DDX3Y* are more dosage-sensitive than other human X-Y gene pairs. For each of the 17 gene pairs, we tallied whether dosage sensitivity had necessitated 1) expression from Xi in human females and 2) maintenance of a Y-homolog in males of diverse species – both features of highly dosage-sensitive genes (Bellott et al. 2014). We addressed the first point by re-analyzing Xi expression data recently generated from cultured human cells (San Roman et al. 2023). We addressed the second point by examining whether the Y-linked gene is conserved across 15 therian (placental mammalian) species where high-quality, contiguous sequence assemblies of the sex chromosomes are available. Specifically, for each Y-homolog, we calculated a phylogenetic branch length – the sum of all branch lengths connecting species where the gene is present, and thus a measure of the gene’s longevity on therian Y chromosomes. We also calculated, for each Y-homolog, the survival fraction across the 15 species – the ratio of observed phylogenetic branch length to maximum possible branch length across the set of species examined (Bellott and Page 2021).

Among X-Y gene pairs, those with the highest dosage sensitivity should be expressed from Xi in females and be long-lived and universally retained on the Y chromosome across species, *i.e.*, have a survival fraction of 1. We find that, among the 17 human X-Y gene pairs, only *DDX3X(Y), KDM6A(UTY)*, and *ZFX(Y)* are expressed from Xi in human females and retain a Y-homolog in all 15 eutherian species examined (Table 1).

**Table 1:**
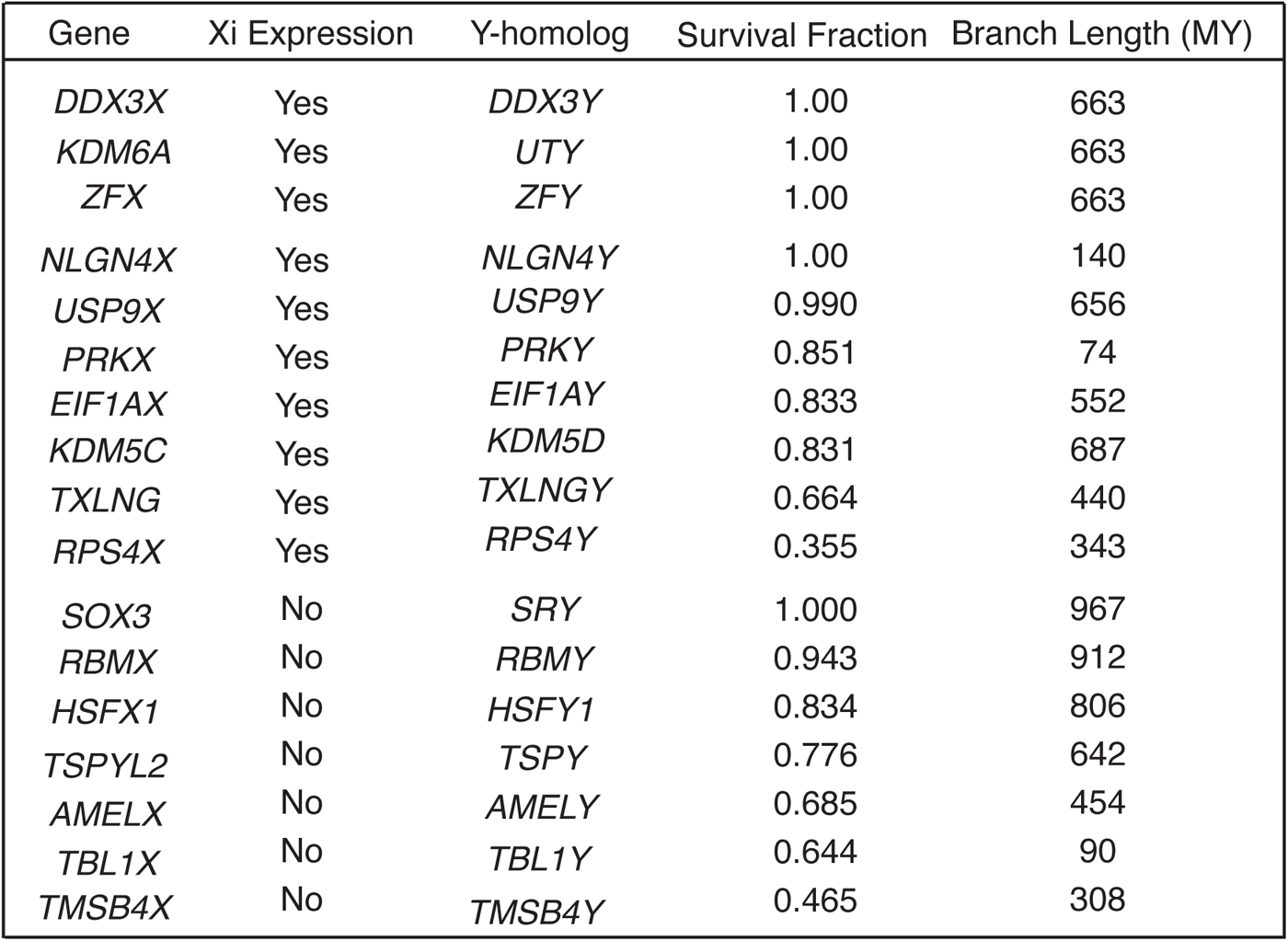
Dosage-sensitivity of human X-Y pair genes across therian mammalian lineages. Xi expression is indicated for X-homologs, and survival fraction and branch length are calculated for the corresponding Y-homologs. Genes are sorted first by Xi expression, then by Y-homolog survival fraction, and finally by Y-homolog branch length.

We further profiled the sensitivity of *DDX3X* to dosage changes using two metrics: 1) P_CT_ scores, which measure the evolutionary conservation of microRNA targeting sites in a gene’s 3’ UTR (Friedman et al. 2009), and 2) LOEUF values, the ratio of observed to expected loss-of-function variants in a gene in human populations (Karczewski et al. 2020) (Supplemental Table S1). High conservation of miRNA targeting sites in a gene’s 3’ UTR implies sensitivity to over-expression (Naqvi et al. 2018), while a low LOEUF value demonstrates sensitivity to diminished function. We rank-ordered all non-PAR genes on the human X chromosome by each of these two metrics (San Roman et al. 2023), from least to most constrained. Among X-Y pair genes expressed from Xi, *DDX3X* has the highest combined sensitivity to over-expression and diminished function, implying that its level of expression is especially constrained (Fig. 1A, Supplemental Table S1).

**Fig. 1:**
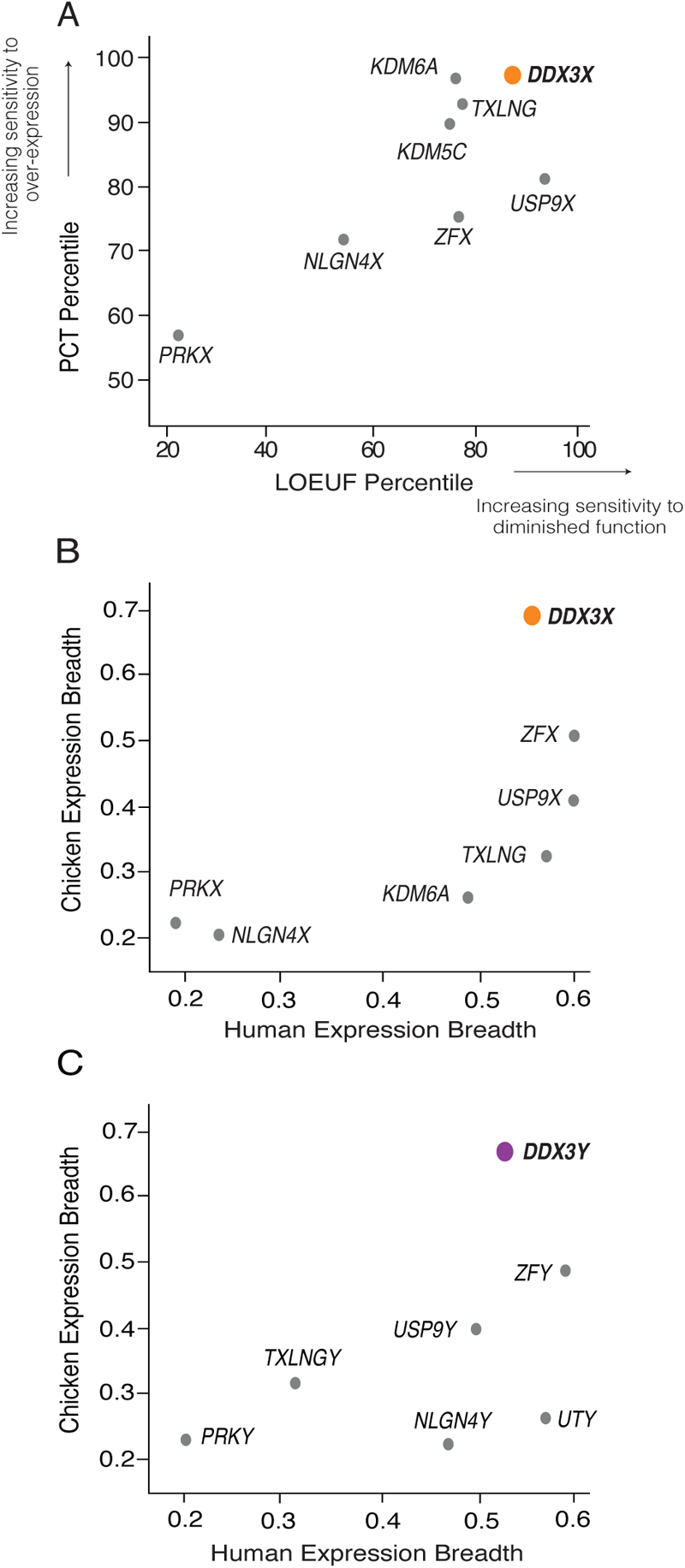
*DDX3X* is highly dosage sensitive and expressed broadly among human tissues. A) Among human X-Y pair genes, *DDX3X* ranks highest in combined sensitivity to over-expression (as judged by PCT percentile among all X-chromosome genes) and diminished function (as judged by LOEUF percentile among all X-chromosome genes). B) *DDX3X* and C) *DDX3Y* and their chicken ortholog display the highest expression breadth among, respectively, the X and Y members of human X-Y gene pairs. Note that expression breadth data was not available for the chicken ortholog of *KDM5C/D*.

We also assessed whether *DDX3X* and *DDX3Y* are expressed more broadly across the body than other X-Y gene pairs – another feature of highly dosage-sensitive genes (Bellott et al. 2014) – and if this breadth was present ancestrally. The ancestral state of sex-linked genes can be inferred from analyses of birds such as chickens, where the orthologs of human sex chromosomal genes are found on autosomes 1 and 4 (Bellott et al. 2010). For each gene pair for which expression data was available in humans (GTEx Consortium, 2017) and chickens (Bellott et al. 2014; Merkin et al. 2012), we measured how broadly the chicken gene and human gene pair were expressed across the body’s various tissues. *DDX3X*, *DDX3Y*, and their autosomal chicken ortholog display the highest combined expression breadth across the two species, suggesting that their dosage is critical throughout the body (Fig. 1B,C, Supplemental Table S2).

### *DDX3X* and *DDX3Y* transcript levels fall as, respectively, Y-chromosome and X-chromosome copy numbers rise

To identify mechanisms that regulate *DDX3X* and *DDX3Y* expression in human cells, we re-analyzed RNA-sequencing data from primary skin fibroblasts of human donors with sex chromosome aneuploidies (San Roman et al. 2023). We first assessed *DDX3X* and *DDX3Y* transcript levels in cells with a single X chromosome and increasing numbers of Y chromosomes (Supplemental Table S3). As expected, *DDX3Y* transcript levels rise with increasing numbers of Y chromosomes. However, *DDX3X* expression from the single X chromosome falls significantly (Fig. 2A,B). Conversely, in cells with a single Y chromosome and increasing numbers of X chromosomes, *DDX3X* transcript levels rise, as expected given the gene’s expression from both Xa and Xi. However, *DDX3Y* expression from the single Y chromosome falls significantly (Fig. 2C,D).

**Fig. 2:**
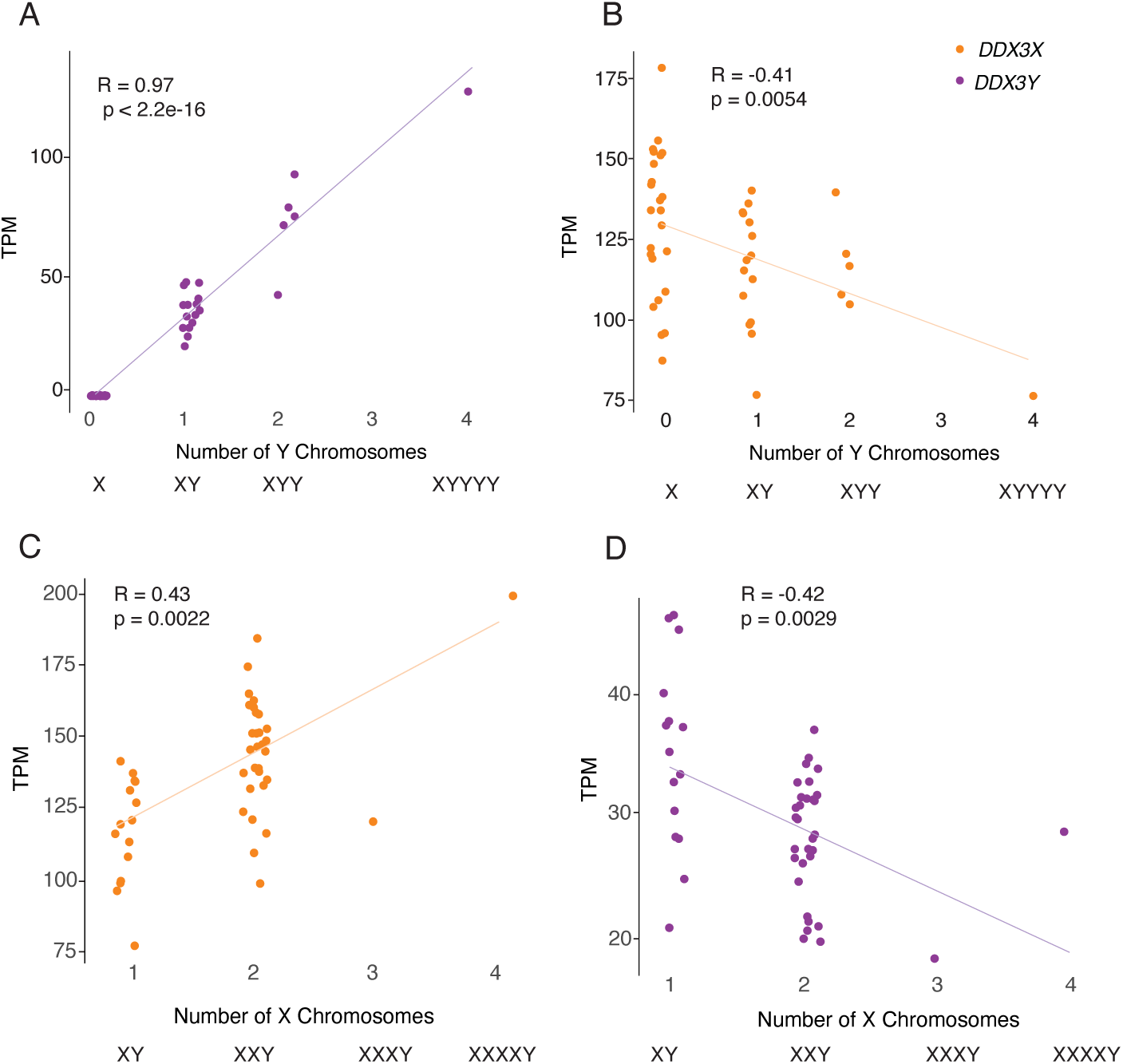
*DDX3X* and *DDX3Y* transcript levels are negatively responsive to, respectively, Y and X chromosome copy number. Scatterplots show *DDX3X* and *DDX3Y* transcript levels in cultured fibroblasts with the indicated sex chromosome constitutions. Each point represents a primary fibroblast culture from one individual. A,B) *DDX3Y* transcript levels are significantly elevated and *DDX3X* transcript levels significantly reduced in fibroblasts with multiple Y chromosomes. C,D) *DDX3X* transcript levels are significantly elevated and *DDX3Y* transcript levels significantly reduced in fibroblasts with multiple X chromosomes. R values and statistical significance calculated using Pearson correlation.

We asked whether this inverse relationship is shared across all X-Y gene pairs or is a unique feature of *DDX3X* and *DDX3Y*. For each X-Y pair gene, we obtained values for the change in its transcript levels per added Xi, and the change in its transcript levels per added Y chromosome (San Roman et al. 2023). In both fibroblasts and lymphoblastoid cell lines (LCLs), *DDX3X* transcript levels fall significantly as the Y chromosome copy number increases; conversely, *DDX3Y* transcript levels fall as the X chromosome copy number increases (Supplemental Table S3). This response is not observed with other X-Y pair genes; it is unique to *DDX3X* and *DDX3Y* (Supplemental Table S3).

We considered the possibility that these decreases in *DDX3X* and *DDX3Y* transcript levels in response to changes in sex chromosome copy number might reflect a general cellular response to aneuploidy. To test this, we examined data from individuals with trisomy 21 (San Roman et al 2023). We observed no change in *DDX3X* or *DDX3Y* transcript levels in response to chromosome 21 copy number (Supplemental Fig. S1). We conclude that *DDX3X* and *DDX3Y* transcript levels are inversely related to Chr Y and Chr X copy numbers, respectively.

### Perturbing *DDX3X* elicits an opposing response in *DDX3Y*, and vice versa

We asked whether these effects of altering sex chromosome copy number are due to *DDX3X* and *DDX3Y* expression changes. We profiled cells with naturally occurring mutations that affect *DDX3X* or *DDX3Y* expression and performed experimental knockdowns to capture the effects of perturbing *DDX3X* and *DDX3Y* transcript levels (Supplemental Tables S4-9).

First, we quantified *DDX3X* transcripts in LCLs from azoospermic (infertile) males with *AZFa* micro-deletions. *AZFa* micro-deletions result from homologous recombination between endogenous retroviral elements on the human Y chromosome, and they remove the *DDX3Y* and *USP9Y* genes without affecting other genes (Fig. 3A) (Sun et al. 2000). We found that *DDX3X* transcript levels were significantly higher in LCLs from *AZFa*-deleted males compared to males with intact Y chromosomes (Fig. 3B, Supplemental Table S4). To test whether *DDX3X* transcript levels are elevated upon deletion of other Y-chromosome regions, we analyzed data from XY individuals whose Y-chromosomes retain *DDX3Y* but are missing several other genes, including the sex-determining gene *SRY* (Schiebel et al. 1997). *DDX3X* transcript levels were unaltered in these individuals (Supplemental Fig. S2, Supplemental Table S5, San Roman et al 2023), demonstrating that *DDX3X* levels are specifically elevated in response to *DDX3Y* deletion.

**Fig. 3:**
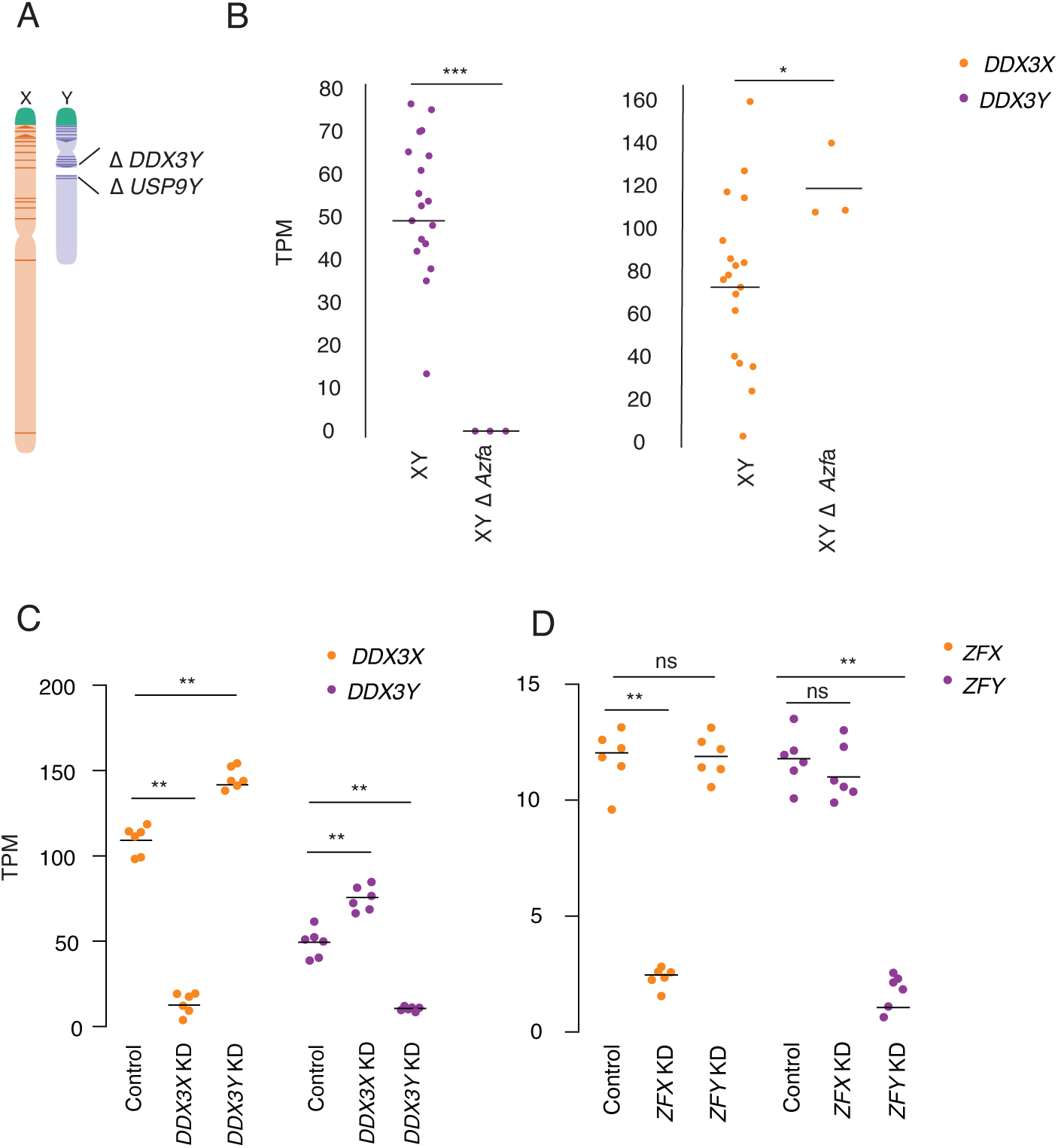
*DDX3X* and *DDX3Y* each respond to perturbations in the other’s expression. A) Schematic diagram of naturally occurring human Y-chromosome (*AZFa*) micro-deletion of *DDX3Y* and *USP9Y*. B) *DDX3X* transcript levels are significantly higher in *AZFa*-deleted 46,XY LCLs compared to Y-chromosome-intact 46,XY LCLs. Each point represents a sample from one individual. Statistical significance determined by Mann Whitney-U test, *** p < 0.0001, * p< 0.05 C) CRISPRi-mediated knockdown of *DDX3Y* using two independent gRNAs in three unrelated 46,XY fibroblast cultures results in significantly elevated *DDX3X* transcript levels. Conversely, *DDX3X* knockdown results in significantly elevated *DDX3Y* transcript levels. D) Re-analysis of CRISPRi knockdown of *ZFX* or *ZFY* (San Roman et al. 2024) demonstrates that knockdown of either gene does not result in significant elevation of the homolog’s transcripts. Statistical significance determined by ANOVA, ** p < 0.001.

We then used CRISPRi to target *DDX3X* or *DDX3Y* for knockdown in primary 46,XY fibroblasts. *DDX3X* transcript levels rose significantly upon knockdown of *DDX3Y* (*DDX3Y* KD), and *DDX3Y* transcript levels responded in a reciprocal fashion to *DDX3X* KD (Fig. 3C, Supplemental Table S6). This negative cross-regulation across X and Y homologs was specific to *DDX3X* and *DDX3Y*; data from CRISPRi knockdowns of *ZFX* and *ZFY*, another broadly expressed, dosage-sensitive X-Y gene pair, did not show this pattern (Fig. 3D, Supplemental Table S7) (San Roman et al. 2024). We validated these findings in an independent dataset, the Cancer Cell Line Encyclopedia (CCLE), which catalogs mutational and expression data from hundreds of cancer cell lines (Ghandi et al. 2019). There we identified 491 different XY cell lines that retained the Y chromosome, and among these, a set of 11 lines that harbored loss of function mutations in *DDX3X* (Supplemental Table S8). *DDX3Y* transcript levels are significantly higher in these 11 cell lines compared to lines where *DDX3X* is intact (Supplemental Fig. S3, Supplemental Table S9). Thus, knockdowns or loss of function in either *DDX3X* or *DDX3Y* are consistently buffered by compensatory increases in the homolog’s expression, demonstrating that *DDX3X* and *DDX3Y* are negatively cross-regulated.

### Negative cross-regulation of *DDX3X* buffers total levels of *DDX3X* and *DDX3Y*

We hypothesized that negative cross-regulation of *DDX3X* and *DDX3Y* maintains the combined expression of the two genes in a narrow range, buffering total transcript levels against changes in gene dosage. To test this, we summed transcript levels for the two genes in our knockdown models. We observed that, in the setting of *DDX3Y* knockdown, the increase in *DDX3X* transcript levels fully compensates and maintains the summed transcript levels of *DDX3X* and *DDX3Y* at control levels (Fig. 4A, Supplemental Table S6). However, in the setting of *DDX3X* knockdown – a larger perturbation – the increase in *DDX3Y* transcript levels does not fully compensate.

We confirmed these results at the protein level using a mass-spectrometry framework that enables sensitive protein quantification by multiplexing peptides and samples (Derks et al. 2022). To measure the summed expression of DDX3X and DDX3Y protein, we quantified peptides shared by DDX3X and DDX3Y (Fig. 4B, Supplemental Table S10).

**Fig. 4:**
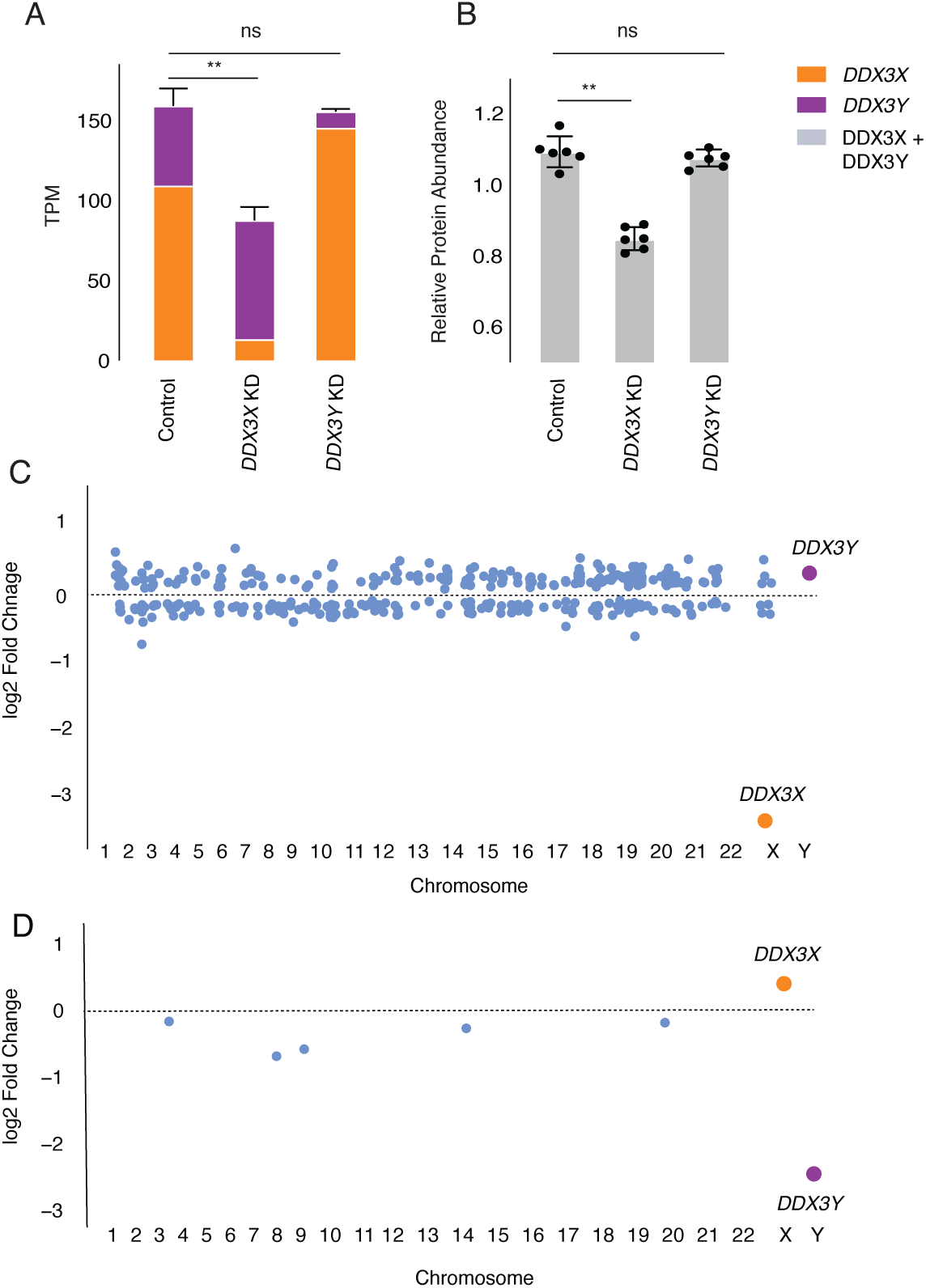
Increased expression of *DDX3X* fully compensates, at transcript and protein levels, for CRISPRi knockdown of *DDX3Y*, but the inverse is not true. A) Stacked bar graph showing summed TPM of *DDX3X* and *DDX3Y* transcripts in knockdowns using two independent gRNAs in three independent 46,XY fibroblast cultures. Statistical significance calculated by ANOVA, ** p < 0.001. B) Bar graph showing abundance of shared DDX3X and DDX3Y peptides in CRISPRi knockdowns with three technical replicates in two independent 46,XY fibroblast cultures. C) Differential gene expression analysis of control vs *DDX3X* knockdown reveals significant expression changes in 397 target genes across the genome, including *DDX3X*. Genes with p < 0.05 (after multiple hypothesis correction) indicated in blue, with exception of *DDX3X* (in orange) and *DDX3Y* (in purple). D) Differential gene expression analysis of control vs *DDX3Y* knockdown reveals only six genes, including *DDX3Y*, that change significantly.

Given these results, we predicted that the incomplete compensation of summed DDX3X + DDX3Y protein levels seen with the *DDX3X* KD would result in transcriptome-wide changes, while such changes would not occur with the *DDX3Y* KD. Indeed, the *DDX3X* KD significantly altered the expression of 379 genes (Fig. 4C, Supplemental Table S11). By contrast, the *DDX3Y* KD significantly altered the expression of only six genes genome-wide, indicating nearly complete compensation through elevated *DDX3X* expression (Fig. 4D, Supplemental Table S12). The *DDX3X* knockdown has far-reaching consequences because of the limited ability of *DDX3Y* to compensate for diminished *DDX3X* expression.

We then asked whether negative cross-regulation of *DDX3X* and *DDX3Y* dampens differences in genome-wide gene expression that might otherwise be observed in individuals with sex chromosome aneuploidies. We found no significant overlap between 1) the set of genes differentially expressed in our *DDX3X* KD and 2) the set of genes transcriptionally responsive to increasing numbers of X chromosomes in the aneuploidy dataset (San Roman et al. 2024) (Fig. S4A). We conclude that, unlike *ZFX*, which drives a large portion of the genome-wide response to X-chromosome copy number (San Roman et al. 2024), *DDX3X* expression that is elevated upon addition of Xi does not drive significant gene expression changes in the aneuploid lines. Indeed, the increase in summed *DDX3X* and *DDX3Y* transcript levels per additional sex chromosome (X or Y) is more modest than that of similarly constrained X-Y pairs (Supplemental Fig. S4B,C,D), consistent with the concept that *DDX3X* and *DDX3Y* are not prominent drivers of gene expression differences associated with sex chromosome aneuploidy.

### *DDX3X* is negatively auto-regulated in 46,XX cells

We hypothesized that negative cross-regulation of the *DDX3X*-*DDX3Y* gene pair evolved from an earlier system of negative auto-regulation in the autosomal ancestor of this X-Y pair. Indeed, *Ded1*, the yeast ortholog of *DDX3X*, appears to be negatively auto-regulated (Silvia Marina et al. 2015). If negative cross-regulation in human XY cells evolved from negative auto-regulation, we might expect to observe negative auto-regulation of *DDX3X* in human 46,XX cells. We set out to test for this and, if present, to ask whether it might be unique among the 17 human NPX genes with NPY homologs. For each X-Y pair gene where informative SNPs could be identified, we obtained its allelic ratio (AR), the ratio of Xi- and Xa-derived transcripts (San Roman et al. 2023). For each gene, we then compared its AR value to its ΔE_X_ value, the increment of change in a gene’s expression per additional X, relative to Xa (San Roman et al. 2023). If an X-linked gene’s expression from Xi and Xa are independent and additive, then the gene’s AR should approximate its ΔE_X_. We found this to be true for other NPX genes with NPY homologs. By contrast, while *DDX3X* has an AR of 0.55 in LCLs and 0.42 in fibroblasts, it has a significantly lower ΔE_X_ of 0.26 in LCLs and 0.16 in fibroblasts (Fig. 5A, Supplemental Table S13). In other words, while Xi contributes 55% or 42% as many *DDX3X* transcripts as Xa does, *DDX3X* transcript levels increase by only 26% or 16% with each additional Xi. In the context of our other findings, this strongly suggests that *DDX3X* is negatively auto-regulated in the absence of *DDX3Y*.

**Fig. 5:**
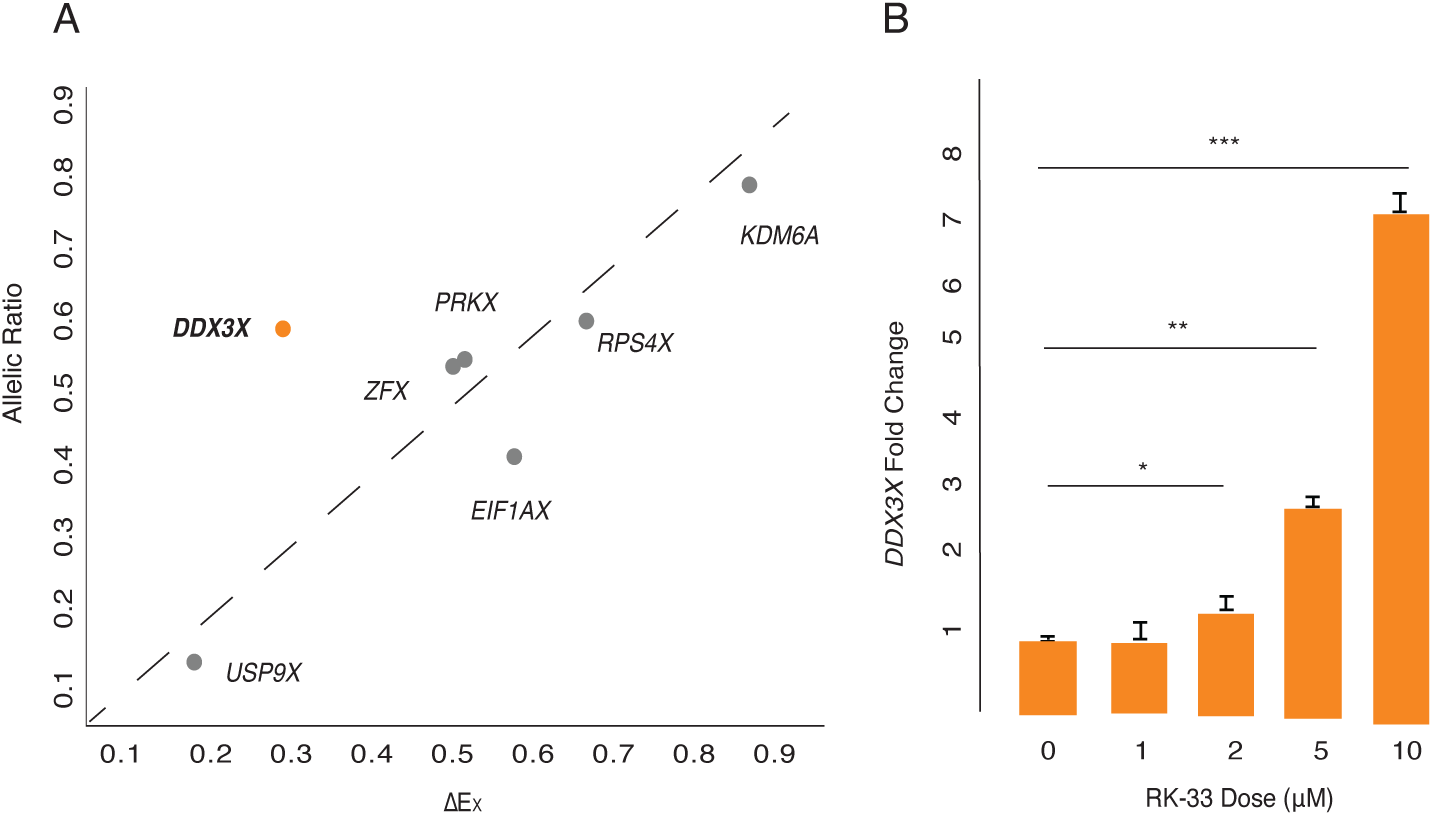
*DDX3X* is negatively auto-regulated in 46,XX cells. A) *DDX3X*’s allelic ratio (AR) is significantly higher than its ΔE_X_ value in LCLs, setting it apart from all other Xi/Xa/Y-expressed X-Y pair genes, whose AR values approximate their ΔE_X_ values. Statistical significance determined via one sample t-test, p = 0.02. B) *DDX3X* transcript levels (by qPCR) in 46,XX fibroblasts are significantly elevated in a dose-responsive manner upon treatment with DDX3 inhibitor RK-33. Statistical significance determined by one-sided t-test on delta Ct values. Error bars indicate standard deviation of three technical replicates. * p < 0.05, *** p < 0.001.

We also hypothesized that chemical inhibition of DDX3X protein activity could lead to increased *DDX3X* transcript levels. To test this, we treated 46,XX fibroblasts with RK-33, a competitive inhibitor of DDX3X that occupies its ATP-binding cleft and disrupts helicase function (Bol et al. 2015). *DDX3X* transcript levels were significantly elevated, in a dose-dependent manner, in cells treated with RK-33, consistent with negative auto-regulation of *DDX3X* in 46,XX cells (Fig. 5B, Supplementary Table S14). Increasing duration of RK-33 treatment also increased *DDX3X* transcript levels in a time-dependent manner (Supplemental Fig. S5, Supplementary Table S15). In summary, allele-specific analysis of transcription and chemical inhibition of protein activity combine to provide evidence for and mechanistic insight into negative auto-regulation of *DDX3X*.

In theory, our observations concerning auto- and cross-regulation could be explained by independent, parallel evolution of negative cross-regulation of *DDX3Y* by *DDX3X*, and of *DDX3X* by *DDX3Y*, but such convergence seems unlikely, especially given the absence of crossing-over as an evolutionary enabler in the case of *DDX3Y*. A simpler hypothesis is that reciprocal cross-regulation of *DDX3X* and *DDX3Y* derives directly from a post-transcriptional mechanism that negatively auto-regulated the ancestral (autosomal) *DDX3* gene. We suggest that this regulatory scheme governed the *DDX3* gene in our amniote ancestors before the autosome carrying *DDX3* became part of today’s (eutherian) mammalian sex chromosomes.

### *DDX3X* response is mediated by mRNA stability

*DDX3X* encodes an RNA-binding protein known to bind its own transcripts (Van Nostrand et al. 2020). In yeast, the *DDX3* ortholog *Ded1* is negatively auto-regulated, and this regulation is dependent on its 3’ UTR (Silvia Marina et al. 2015), indicating that *Ded1* mRNA stability is being modulated. We reasoned that the negative cross-regulation we observed between human *DDX3X* and *DDX3Y* may also involve mRNA stability. If *DDX3Y* destabilizes *DDX3X* transcripts, we would expect the half-life of *DDX3X* transcripts to decrease in response to increasing *DDX3Y* dosage. We tested this prediction by labeling nascent mRNAs in 46,XY and 49,XYYYY LCLs with 5-EU and sequencing the resultant mRNA populations at discrete intervals to quantify half-life (Fig. 6A). We calculated the ratio of nascent mRNA/total mRNA normalized to steady-state levels across time points, and we observed a striking difference in *DDX3X* mRNA half-life between the two conditions. *DDX3X* mRNAs have a half-life of 0.5h in 49,XYYYY cells compared to 1.3h in XY cells (Fig. 6B, Supplemental Table S16), implying that high DDX3Y levels lead to a marked destabilization of *DDX3X* mRNAs, reducing steady state levels of *DDX3X* transcripts. This finding was corroborated in an independent metabolic labeling trial with a shorter timecourse (Supplemental Fig. S6A, Supplemental Table S17). As predicted, steady-state levels of *DDX3X* mRNA were lower in 49,XYYYY as compared with 46,XY samples (Supplemental Fig. S6B).

**Fig. 6:**
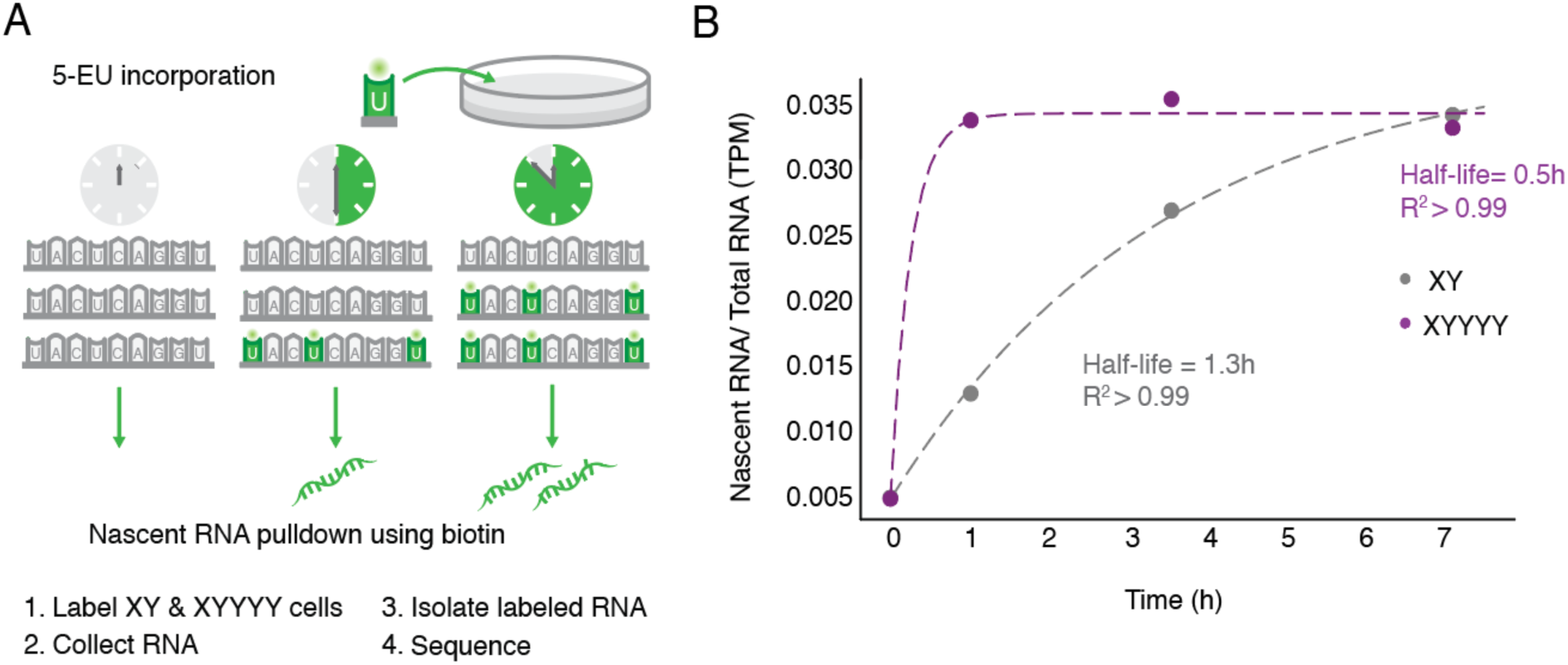
*DDX3X* mRNA stability is regulated. A) Schematic of experiment to determine half-lives of mRNAs. 46,XY and 49,XYYYY LCLs were incubated with 5-ethyl uridine (5-EU) to obtain nascent mRNAs. B) *DDX3X* has an mRNA half-life of 0.5h in 49,XYYYY vs 1.3h in 46,XY LCLs.

These results support a model where the ancestral (autosomal) *DDX3* gene in amniotes destabilized its own transcripts to negatively auto-regulate its expression, foreshadowing the ability of mammalian *DDX3X* and *DDX3Y* to destabilize their own and each other’s transcripts.

**Fig. 7:**
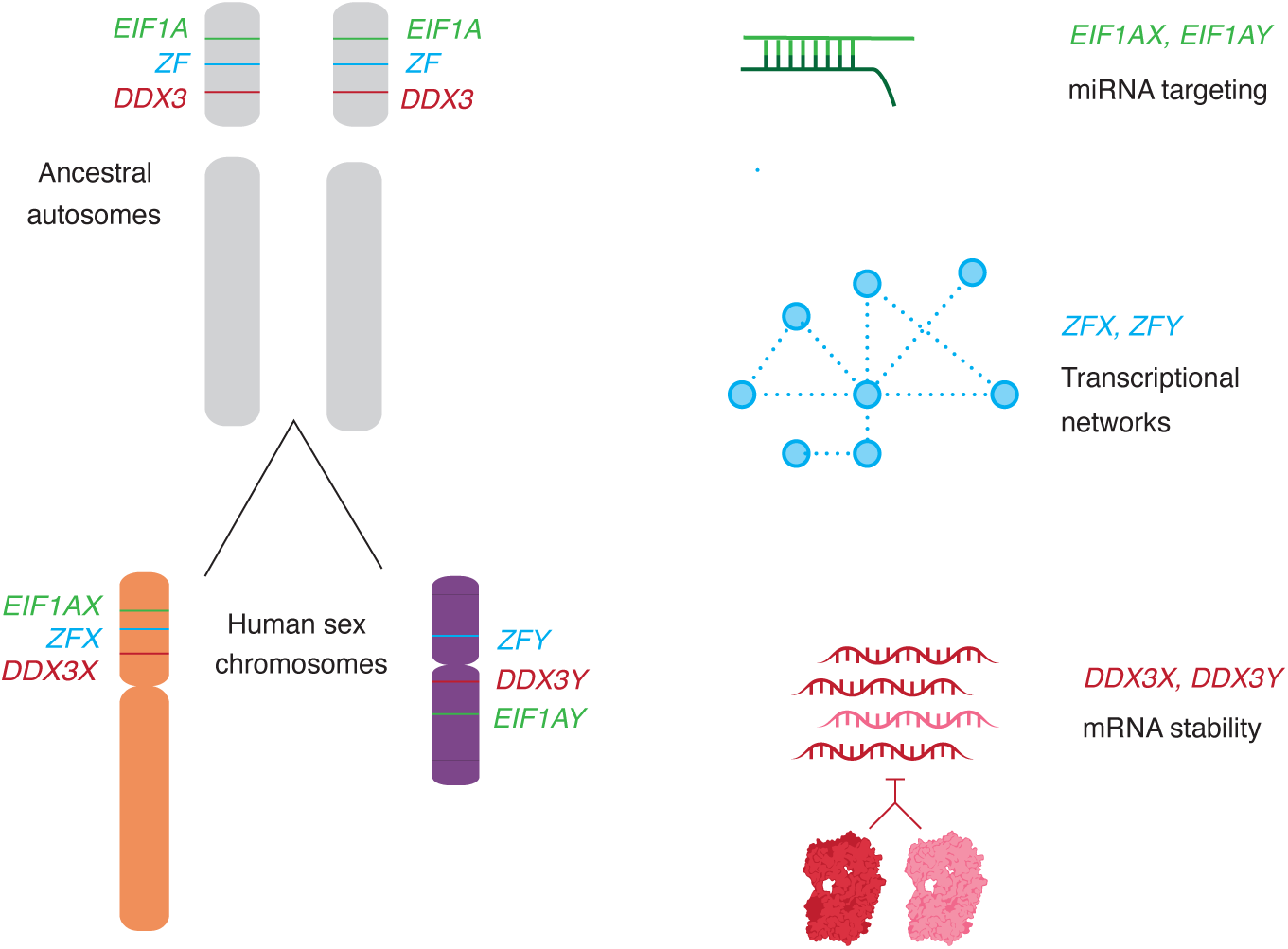
Not only protein-coding sequences, but also gene regulatory mechanisms were preserved during the evolution of sex chromosomes from ordinary autosomes. The auto- and cross-regulation of *DDX3X* and *DDX3Y* reported here likely originated from the auto-regulation of ancestral (autosomal) *DDX3*. Together with published studies of two other X-Y gene pairs -- *EIF1AX-EIF1AY* and *ZFX-ZFY* (Godfrey et al 2020, San Roman et al 2024) – our findings suggest that an array of gene-specific regulatory schemes operative on the ancestral autosomes persist today on the human X- and Y-chromosomes.

## Discussion

As described here, *DDX3X* and *DDX3Y* are negatively, post-transcriptionally cross-regulated (Fig. 3), and *DDX3X* is negatively, post-transcriptionally auto-regulated (Fig. 5), such that perturbations to one allele (of *DDX3X* or *DDX3Y*) can be buffered by upregulation of the other allele. This is the first observation of an X-Y gene pair having cross-regulatory capabilities, and it helps explain certain human phenotypes associated with loss-of-function mutations as well as diverse observations in the literature.

In 46,XY males, rare constitutional (germline) mutations in *DDX3X* cause a neurodevelopmental disorder (‘*DDX3X* syndrome’) (Kellaris et al. 2018; Nicola et al. 2019). By contrast, constitutional mutations of *DDX3Y* in 46,XY males cause a surprisingly subtle phenotype. De novo deletions of the entire *DDX3Y* gene (so-called *AZFa* deletions) cause spermatogenic failure and, thereby, infertility but otherwise have no reported impact on somatic development, function, or health (Fig. 3) (Sun et al. 2000). *In vitro*, in LCLs, we find that elevated *DDX3X* transcript levels compensate for the absence of *DDX3Y* (Fig. 4A,B). We propose that the same holds in the brain (and other somatic tissues) of *AZFa*-deleted males, explaining why males with germline *DDX3Y* deletions display no neurodevelopmental consequences.

Our data indicates that negative cross-regulation of *DDX3X* and *DDX3Y* operates broadly and potentially universally across human somatic cell types. We observe this phenomenon in multiple human cell types: in LCLs, in primary fibroblasts, and in cancer cell lines originating from five different tissues (Fig. 3, Supplemental Table S2, Supplemental Fig. S4). The generality of these findings allows us to reinterpret and better comprehend diverse observations regarding *DDX3X* and *DDX3Y* reported in the literature. Negative posttranscriptional regulation offers a unifying explanation for the following observations: 1) In *DDX3X*-mutant lymphomas in human males, Gong and colleagues reported that *DDX3Y* transcript levels were elevated compared to wild-type lymphocytes (B cells). Gong et al. speculated that *DDX3Y* up-regulation in these *DDX3X*-mutant lymphomas reflected an aberrant, oncogenically adaptive gene expression program (Gong et al. 2021). A simpler explanation is provided by negative cross-regulation that operates universally in human somatic cell types, including cancers of somatic origin. 2) In the brains of male mice bearing various conditional *Ddx3x* knockouts designed to model either human *DDX3X* syndrome (Hoye et al. 2022) or medulloblastoma (Patmore et al. 2020), investigators noted that *Ddx3y* transcript levels were elevated compared to wild-type. Viewed in light of our current findings, these observations suggest that negative cross-regulation of *DDX3X* and *DDX3Y* occurs not only in humans but also in mice.

These post-transcriptional regulatory connections between *DDX3X* and *DDX3Y* are unique, to our knowledge, among human X-Y gene pairs. How could such a system evolve? We considered this question in the context of the evolution of the X and Y chromosomes from ordinary pairs of autosomes during the past 200 million years. We reasoned that tight delimiting of *DDX3* gene expression likely pre-dated the divergence of the homologous genes *DDX3X* and *DDX3Y* on the (eutherian) mammalian sex chromosomes, as this would most economically explain the presence of both auto- and cross-regulation of the human genes. Indeed, there is evidence that *Ded1*, the yeast ortholog of *DDX3X,* is also negatively auto-regulated (Silvia Marina, 2015), suggesting that this regulation has been preserved in both the eutherian and yeast lineages during the 1.3 billion years since their divergence (Kumar et al. 2022). We infer that *DDX3* was already highly dosage sensitive when, as a single-copy gene, it resided on an amniote autosome that later gave rise to much of the sex chromosomes of eutherian mammals. *DDX3X* and *DDX3Y* evidently retained this high dosage sensitivity and the associated negative regulatory scheme that had governed their common autosomal ancestor.

Combined with other recent discoveries, our present findings illuminate the breadth and diversity of gene regulatory mechanisms and networks that were selectively preserved as the X and Y chromosomes evolved from ordinary autosomes during the past 200 million years (Fig. 7). For example, our recent studies of the genome-wide consequences of human sex chromosome aneuploidy showed that the X- and Y-linked transcriptional activators ZFX and ZFY modulate expression of large and remarkably similar sets of autosomal genes (San Roman et al. 2024). Given the scale of these gene regulatory networks, their similarity is unlikely to be the result of convergent evolution. A more economical explanation is evolutionary preservation of preexisting gene regulatory networks centered on the single autosomal forebear of the eutherian *ZFX* and *ZFY* genes. Another example involves our recent observation that expression of the Y-linked translation initiation factor EIF1AY is enhanced (relative to its X-linked homolog, EIF1AX) in the human heart (Godfrey et al. 2020). This was explained through our recent discovery of 1) a miR-1 (cardiac microRNA) binding site in the 3’ UTR of the ancestral (autosomal) *EIF1A* gene, 2) preservation of that ancestral binding site in the 3’ UTR of *EIF1AX*, and 3) loss of the ancestral binding site in the 3’ UTR of *EIF1AY*, resulting in its enhanced expression in the human heart. In sum, the cases of *DDX3X*/*Y*, *ZFX*/*Y*, and *EIF1AX*/*Y* illustrate the diversity and reach of ancestral (autosomal) gene regulatory mechanisms preserved, or in some cases lost, during the 200-million-year evolution of the eutherian sex chromosomes from ordinary autosomes.

## Methods

### Analysis of total branch length and survival fraction

For each gene, total branch length and survival fraction values in therian species were obtained from Bellott and Page (Bellott and Page 2021). To obtain a gene’s total branch length, all branch lengths in the most parsimonious tree connecting all species where the gene is present are summed from the last common ancestor. The survival fraction is the observed total branch length divided by the maximum possible branch length. Survival fractions range from 0 (lost in all lineages) to 1 (retained in every lineage).

### Analysis of constraint metrics

We downloaded LOEUF (loss-of-function observed/expected upper fraction) scores from gnomAD (v2.1.1.lof_metris.by_gene.txt;https://gnomad.broadinstitute.org/) and only used scores with a minimum of 10 expected LoF variants. For sensitivity to an increase in gene dosage, we used the per-gene average probability of conserved miRNA targeting scores (PCT) (Friedman et al.2009). We computed a percentile rank score for each metric, from most constrained to least constrained (San Roman et al. 2023). Pythagorean sum of ranks was used to calculate a combined metric for dosage sensitivity (Supplemental Table 1).

### Calculation of expression breadth

Human expression breadth was calculated from GTEx v8 using male samples. For each gene, expression breadth was calculated using TPM values as follows: Sum of expression across tissues/(Maximum expression in a tissue * Number of tissues). For each X-Y gene pair, expression breath values for the X-homolog and Y-homolog were averaged to generate a mean score. Chicken expression breadth values were obtained from Bellott et al using data from Merkin et al. (Bellott et al. 2010; Merkin et al. 2012). Pythagorean sum of breadths was used to calculate a combined metric for dosage sensitivity (Supplemental Table 2).

### Aneuploidy data

RNA-sequencing data from cultured cells of individuals with sex chromosome aneuploidy (San Roman et al. 2023) were downloaded from https://doi.org/10.1016/j.xgen.2023.100259.

### Cell Culture

All LCLs were cultured in complete RPMI at 37C. Fibroblasts were cultured in high-glucose DMEM (Gibco), 20% FBS, L-Glutamine (MP Biomedicals), MEM Non-Essential Amino Acids (Gibco), 100 IU/ml Penicillin/Streptomycin (Lonza).

### CRISPRi

Three independent, unrelated 46,XY fibroblast cultures stably expressing a nuclease-dead Cas9 fused with a repressive KRAB domain (dCas9-KRAB) were obtained from Adrianna

San Roman. gRNAs for control (intergenic), *DDX3X*, and *DDX3Y* were chosen from the human CRISPRi v2 library (Horlbeck et al. 2016) and cloned into the sgOpti lentiviral expression vector. Viral particles were generated and frozen as described in San Roman et al. Guide sequences were as follows:

Control 1: GACATATAAGAGGTTCCCCG

Control 2: AACGGCGGATTGACCGTAAT

*DDX3X* #1: GTCCCGTGAGAGGGCCTTCG

*DDX3X* #2: GCCCGGGACGAGCACAATGG

*DDX3Y* #1: GTTCGGTCTCACACCTACAG

*DDX3Y* #2: GAGTACTGGGCCTCACGCAA

Control and *DDX3X*- or *DDX3Y*-targeting gRNAs were transduced into the stably-expressing dCas9-KRAB fibroblasts, and cells were selected using 2 ug/mL puromycin (Sigma) beginning 24 h post infection. Cells were washed once with PBS and collected 72 h post infection. RNA was extracted with the RNeasy Mini Kit (Qiagen). RNA sequencing libraries were prepared using the KAPA mRNA HyperPrep Kit V2 (Roche). Paired-end 100x100 bp sequencing was performed on a NovaSeq 6000 (Illumina). Reads were pseudoaligned with kallisto and imported into R using tximport. Differential gene expression analysis was performed using DESeq2 (Love et al. 2014). RNA-sequencing data from *ZFX* and *ZFY* knock-down experiments (San Roman et al. 2024) were downloaded from dbGaP Study Accession: phs002481.v2.p1.

### Treatment with RK-33

46,XX fibroblast cultures were treated with 0, 1, 2, 5, or 10 μM RK-33 in DMSO for 24 h. For the time course, they were treated with 2 μM RK-33 for 0, 1, 2, 4, or 24 h.

### qPCR

Cells were washed once with PBS and collected 72 h post treatment. RNA was extracted with the RNeasy Mini Kit (Qiagen) and cDNAs prepared with Super-Script Vilo Master Mix (Thermo Fisher). *DDX3X* levels were quantified by qPCR using Fast Sybr Green Master Mix (Thermo Fisher). Primers for *DDX3X* and reference gene *ACTB* were as follows:

*DDX3X* F: GTGGAAGTGGATCAAGGGGA

*DDX3X* R: TGATTTGTCACACCAGCGAC

*ACTB* F: CACCAACTGGGACGACAT

*ACTB* R: ACAGCCTGGATAGCAACG

### Analysis of Cancer Cell Line Expression dataset

Expression and mutation data for cancer cell lines were downloaded from the DepMap 22Q2 release (https://depmap.org/portal/download/all/). Analysis was restricted to 46,XY cells by applying a log2TPM filter of > 0.2 *DDX3Y,* > 0.2 *RPS4Y*, < 2 *XIST*.

### Sample preparation for mass spectrometry

Samples were prepared for proteomic analysis by mPOP (Minimal Proteomic Sample Preparation) as described in Specht et al (Specht et al. 2018). Briefly, cells were resuspended in MS-grade water and frozen. They were then heated at 90 C for 10 min to lyse cells. Proteins were reduced and treated with Trypsin Gold (Promega). The peptide abundance of each sample was measured and each sample was labeled with non-isobaric mass tags, mTRAQ Δ0, Δ4, or Δ8 (SciEx: 4440015, 4427698, 4427700) following the manufacturer’s instructions. Reactions were quenched and pooled as a 3-plex with relative mass offsets of Δ0, Δ4, and Δ8.

### Mass Spectrometry data acquisition

mTRAQ-labeled peptide sets were separated by reversed-phase UHPLC in 1 µl injections by a Dionex UltiMate 3000 using a 25 cm × 75 µm IonOpticks Aurora Series UHPLC column (AUR2-25075C18A). Buffer A was 0.1% formic acid in MS-grade water. Buffer B was 80% acetonitrile (ACN) with 0.1% formic acid, mixed in MS-grade water. The gradient was as follows: 4% Buffer B (minutes 0–11.5), 4–7% Buffer B (minutes 11.5–12), 7–32% Buffer B (minutes 12–75), 32–95% Buffer B (minutes 75–77), 95% Buffer B (minutes 77–80), 95–4% Buffer B (minutes 80–80.1), and 4% Buffer B until minute 95. The flowrate was 200 nL/min throughout.

Mass spectrometry data were acquired using a DIA method which utilizes frequent MS1-scans for quantitation, as previously described (Derks et al. 2022). The duty cycle consisted of 5 sub-cycles of (1 MS1 full scan × 5 MS2 windows) for a total of 25 MS2 windows to span the full m/z scan range (380–1,370 m/z). MS1 and MS2 scans were performed at 140k and 35k resolving power, respectively.

### Mass spectrometry data analysis

Raw plexDIA data were processed with DIA-NN (version 1.8.1 beta 16) (Demichev et al. 2019) using the following settings and additional commands: {--window 1}, {--mass-acc 10.0}, {--mass-acc-ms1 5}, {--reanalyse}, {--rt-profiling}, {--peak-height}, {--fixed-mod mTRAQ, 140.0949630177, nK}, {--channels mTRAQ,0,nK,0:0; mTRAQ,4,nK,4.0070994:4.0070994; mTRAQ,8,nK,8.0141988132:8.0141988132}, {--peak-translation}, {--original-mods}, {--report-lib-info}, {--ms1-isotope-quant}, {--mass-acc-quant 5.0}.

The resulting data were filtered at 1% FDR for precursors and protein-groups (DDX3X;DDX3Y). Precursors were further filtered for Translated.Q.Value < 0.01. MaxLFQ (Cox et al. 2014) was used to perform protein-group-level quantification for all samples. Each protein-group was normalized to the mean value in each LC-MS run, as each LC-MS run contained three technical replicates from each of the three conditions (control, *DDX3X* knockdown, and *DDX3Y* knockdown). Each sample was then normalized to its own respective median protein group value to account for differences in absolute protein abundances between samples.

Finally, each protein-group was normalized to the mean value of the protein-group across all samples, for each cell-line. For each cell line, batch-correction was performed using Combat (Leek et al. 2012) with missing data imputed with a kNN algorithm (k=3), to correct biases produced by using different mass-tag offsets (e.g. Δ0, Δ4, and Δ8).

### 5-EU labeling and cell collection

LCLs were thawed and allowed to grow in T175 flasks. Cells were split with fresh LCL media and 5EU (Jena Bioscience) was added to a final concentration of 400 µM. Cells were collected 0, 0.5,1, 1.5, and 2, 3.5 and 7 h later, washed with PBS and pelleted prior to addition of TRIzol (Thermo Fisher Scientific) reagent. Cells were snap-frozen at ^-^ 80 C. RNA was precipitated with isopropanol and 1ng of 5-EU EGFP positive control was added.

### Biotinylation and pulldown

Biotinylation and pulldown were performed as described (Kingston and Bartel 2019). Briefly, biotin was attached to metabolically labeled RNAs in a 10 µL reaction protected from light. The reaction was quenched and RNA precipitated. RNA was then incubated with blocked and pre-washed streptavidin bead slurry. Beads were washed once more and RNA was eluted with TCEP (tris(2-carboxyethyl) phosphine) and water. RNA was precipitated and libraries were then prepared using the SMART-Seq v4 Ultra Low Input RNA kit and sequenced on a NovaSeq 6000. Input RNAs were also sequenced to measure total RNA. TPMs were normalized to 5-EU positive EGFP spike-in. Normalized fraction of nascent/total *DDX3X* mRNA was fit to the equation y = α/β * 1-e^β/t^ to obtain β (half-life).

### Statistical Methods

Various statistical tests were used to calculate p values as indicated in the methods section, figure legends, or text, where appropriate. Results were considered statistically significant when p < 0.05 or FDR < 0.05 when multiple hypothesis correction was applied, unless stated otherwise. All statistics were calculated using R software, version 4.2.1 or Prism, version 9.4.1 unless stated otherwise.

## Data and Materials Access

Original code for the analyses in this paper is deposited at https://github.com/shruthi3195/DDX3X_SR_2023. Raw reads for sequencing data were deposited at NCBI and can be accessed at dbGaP Study Accession: phs002481.v3.p1. All cell lines used in this study are listed in Supplementary Table S18.

## Acknowledgements

We thank members of the Page lab for advice; J. Adarme and S. Tocio for laboratory support; A.K. San Roman and J. Hughes for comments on the manuscript; the Whitehead Institute Genome Technology Core for library preparation and sequencing; and Caitlin Rausch for illustration. Supported by the Howard Hughes Medical Institute, the Whitehead Institute, the Brit Jepson d’Arbeloff Center for Women’s Health, the National Institutes of Health (R35GM148218 to N.S.), and philanthropic gifts from Arthur W. and Carol Tobin Brill, Matthew Brill, Charles Ellis, the Barakett Foundation, the Howard P. Colhoun Foundation, and the Seedling Foundation.

